# Active learning for medium optimization for selective bacterial culture

**DOI:** 10.1101/2023.11.15.567258

**Authors:** Shuyang Zhang, Honoka Aida, Bei-Wen Ying

## Abstract

Medium optimization and development for selective bacterial culture are essential for isolating and functionalizing individual bacteria in microbial communities; nevertheless, it remains challenging due to the unknown mechanisms between bacterial growth and medium components. The present study first tried combining machine learning (ML) with active learning to finetune the medium components for the selective culture of two divergent bacteria, i.e., *Lactobacillus plantarum* and *Escherichia coli*. ML models considering multiple growth parameters of the two bacterial strains were constructed to predict the finetuned medium combinations for higher specificity of bacterial growth. The growth parameters were designed as the exponential growth rate (*r*) and maximal growth yield (*K*), which were calculated according to the growth curves. The eleven chemical components in the commercially available medium MRS were subjected to medium optimization and specialization. High-throughput growth assays of both strains grown separately were performed to obtain thousands of growth curves in more than one hundred medium combinations, and the resultant datasets linking the growth parameters to the medium combinations were used for the ML training. Repeated rounds of active learning (i.e., ML model construction, medium prediction, and experimental verification) successfully improved the specific growth of a single strain out of the two. Both *r* and *K* showed maximized differentiation between the two strains. Further analysis of all data accumulated in active learning identified the decision-making medium components for growth specificity and the differentiated determinative manner of growth decision of the two strains. In summary, this study demonstrated the efficiency and practicality of active learning in medium optimization for selective culture and offered novel insights into the contribution of the chemical components to specific bacterial growth.

## Introduction

Culturomics has emerged as a vital method for studying complex microbial environments. It often combines various medium conditions for selective cultures to identify bacterial species. In environmental microbiology, culturomics has led to a reevaluation of microbial diversity, particularly for those microbes that are challenging to culture ^1^. In clinical microbiology, culturomics has led to the cultivation of 341 bacterial species from 212 different culture conditions, with over half of these being newly discovered in the human gut ^2^. The primary objective in the development of culturomics is to enable the method to provide diverse culture conditions that promote the growth of fastidious bacteria ^3^. With the aim of screening and identifying specific microorganisms within samples, the development of culture media for specific bacterial growth has become increasingly crucial. By incorporating various growth inhibitors into the culture media, unwanted microbial populations can be eliminated, facilitating the growth of the target microorganisms ^4^. Scientists have been exploring novel compositions for culture media, such as those that mimic natural marine environments, leading to the detection of new microorganisms ^5^. Due to the complexity of increasing samples and the demands for screening and identification, medium development for selective culture has faced new challenges ^6, 7, 8^. The selective culture ensured the target bacterial growth and prevented other microbial communities from growing ^4^. The typical approach of adding inhibitors might also suppress the growth of the target bacterium. In food industries, selective culture media were frequently used to detect microbial contamination and spoilage in food materials, which might be unsuitable for competitive bacteria ^9, 10^. Therefore, medium optimization and specialization are highly required in the field.

Medium optimization was challenging due to the high complexity of the microbiomes and combinations of medium components ^11^. Traditional methods of Design of Experiments (DOE) ^12^ and Response Surface Methodology (RSM) ^13, 14^ employed a quadratic polynomial approximation; thus, they might not fully capture the complex interactions between the medium and cells ^15^. Machine learning (ML) has been introduced to predict unknown events by learning a dataset ^16^. This approach has been widely applied in drug development ^17^, protein structure and function prediction ^18^, and epidemic surveillance ^19^, and exhibited better outcomes than DOE or RSM ^20^. Lately, combining active learning with ML has successfully optimized the culture media for mammalian cells ^21^. These studies strongly suggested the efficiency of ML-associated active learning for medium development and its availability to improve the selective effect of culture medium for specific bacterial growth, so-called medium specialization.

In the present study, ML-combined active learning considering single or multiple growth parameters was conducted to finetune the medium compositions for the specific growth of *Lactobacillus plantarum* or *Escherichia coli*. High-throughput growth assays were performed to acquire the training data and for experimental verification. Multiple benchmarks, i.e., scores, were newly designed to associate with ML to predict better medium combinations for specific bacterial growth. The datasets connecting the medium combinations with the goodness of bacterial growth obtained during active learning were analyzed to discover the contribution of medium components to bacterial growth specificity. The decision-making elements for bacterial growth and growth specificity were identified. The study tried to provide a representative case of employing active learning for medium specialization and insights into the medium contribution to selective bacterial culture.

## Results

### Experimental and computational design of the active learning for medium optimization

The initial training data was experimentally acquired, linking medium combinations to bacterial growth. *Escherichia coli* (*Ec*) and *Lactobacillus plantarum* (*Lp*) were used, as they were of different growth preferences and commonly employed in laboratories and production tests using selective culture media ^22, 23, 24, 25^. Although the media appropriate for both strains were well-known, whether the culture medium specific for *Lp* growth could be finetuned via machine learning for *Ec* growth was tested. Eleven components in the commercially available MRS medium for *Lp* growth were used to prepare the medium combinations. These components were mixed in a broad range of concentration gradients, changing in a logarithmic scale (Fig. 1A). *Ec* and *Lp* were cultured independently in 98 medium combinations (N=4) to obtain the growth curves for calculating the growth parameters of growth rate (*r*) and maximal population density (*K*) (Fig. 1B). As the initial training data, the medium combinations connecting with the growth parameters of both strains, i.e., *r_Lp*, *K_Lp*, *r_Ec*, *K_Ec*, were acquired. These four parameters were used as the machine learning (ML) objective variables, either in a single mode or a multiple combination (Fig. 2A).

**Figure 1.**
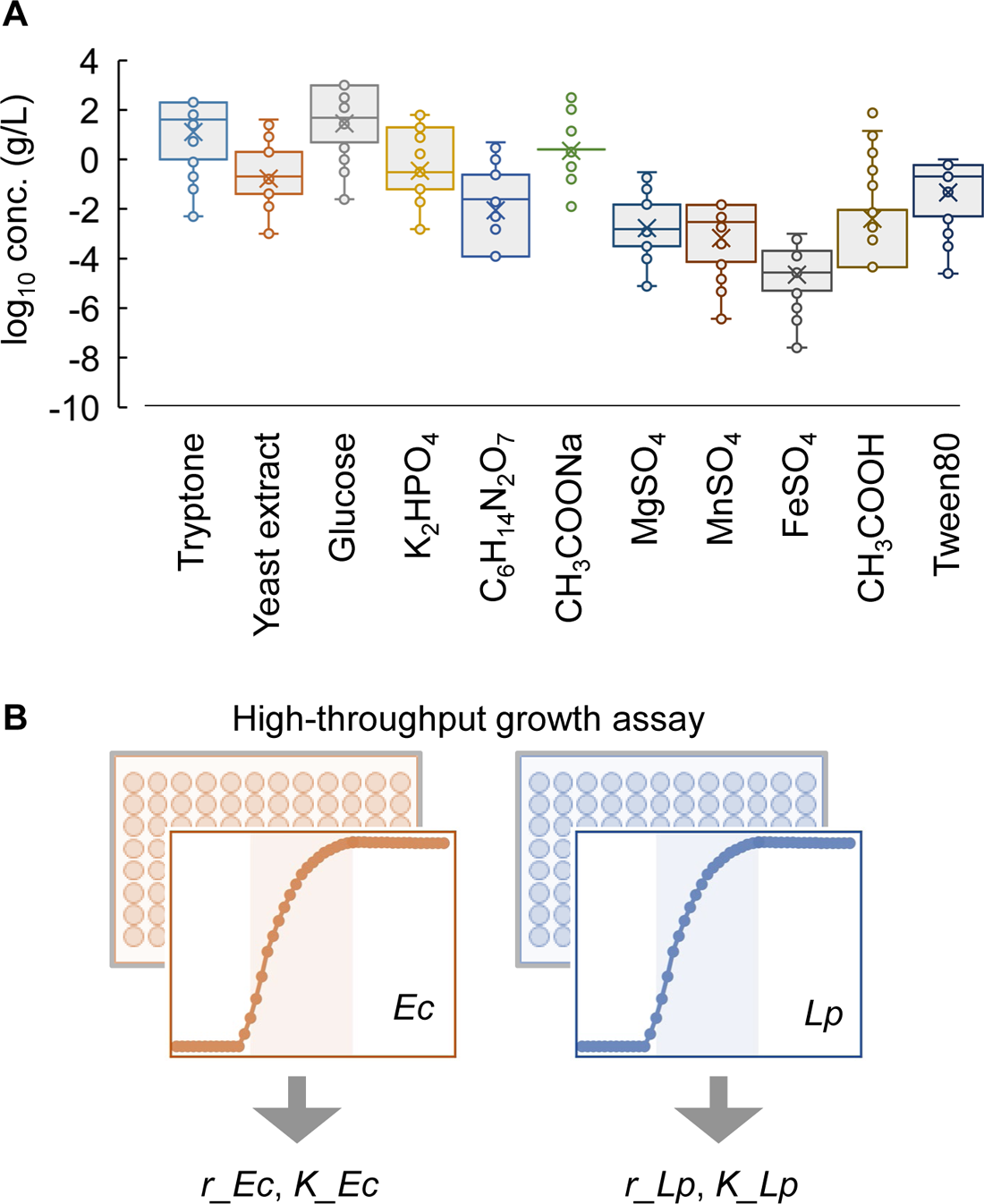
Growth assay under medium combinations. **A.** Concentration gradients of medium components. Circles indicate the concentrations used in the medium combinations, shown on a logarithmic scale. **B.** High-throughput growth assays. Mono-culture of two bacterial strains was performed under hundreds of medium combinations. The growth parameters calculated from the growth curves, i.e., growth rate and growth yield, are indicated as *r* and *K*, respectively.

**Figure 2.**
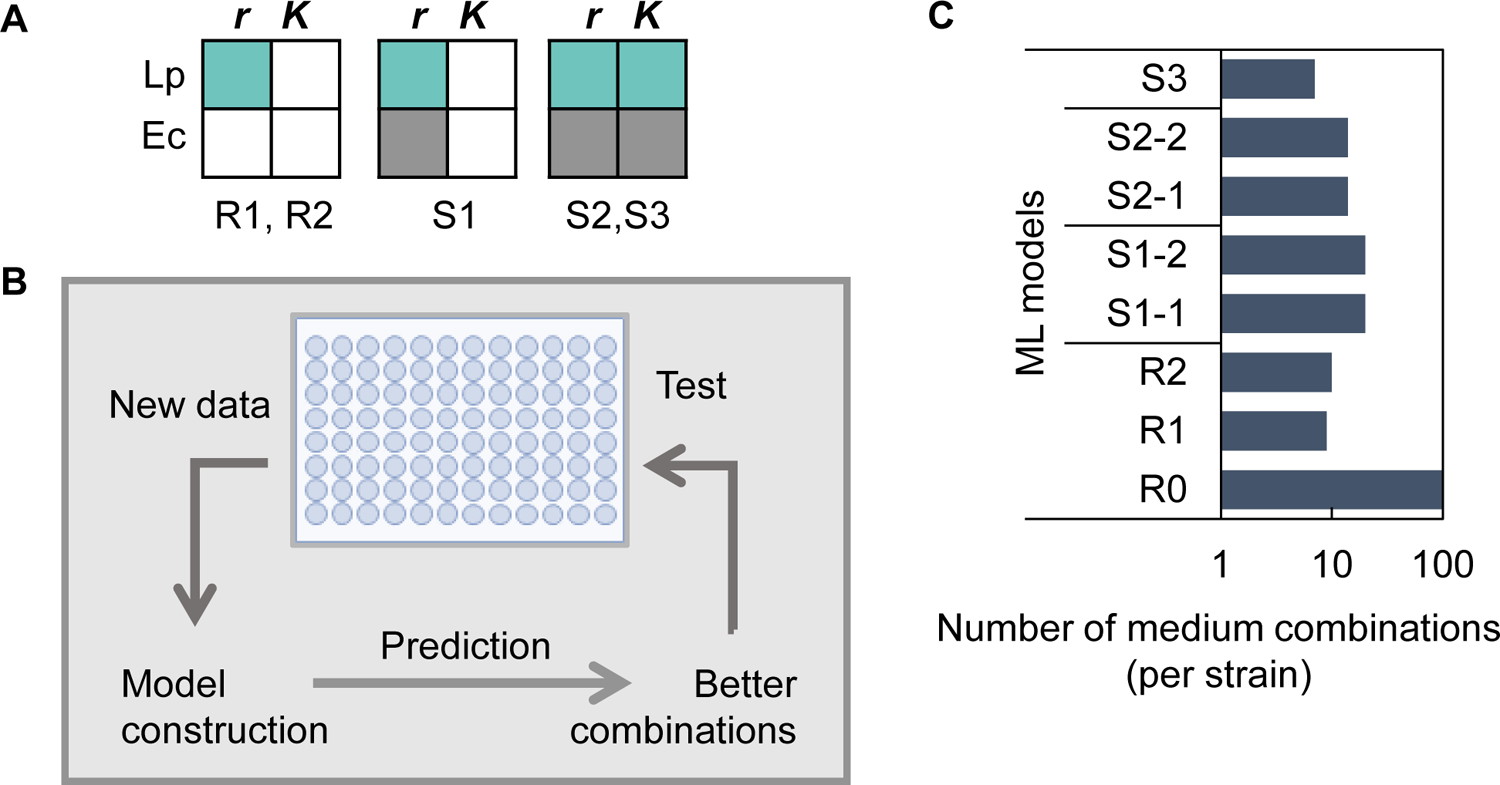
Active learning for medium optimization. **A.** ML models considering single or multiple growth parameters. The growth parameters subjected to be increased or suppressed are indicated in cyan and grey, respectively. R1 and R2 consider a single out of four parameters; S1 considers the paired parameters; S2 and S3 consider all four. **B.** Repeated rounds of active learning. The process of active learning is presented, i.e., ML model construction (described in **A**), medium prediction, and experimental verification. **C.** Number of experimentally tested medium combinations in each round of active learning. R0 indicates the initial training data. S1-1, S1-2, S2-1, and S2-2 represent two rounds of active learning with the ML models of S1 and S2, respectively.

ML-assisted medium optimization and specialization for different strains were performed using the gradient-boosting decision tree (GBDT), which has been repeatedly validated to have superior predictive performance and interpretability compared to other algorithms ^26, 27, 28^. The initial training data (R0) was applied to the GBDT model to improve *r_Lp* or *K_Lp* (R1 and R2) (Fig. 2A). Medium optimization and specialization were performed by active learning, following model construction, prediction, and experimental verification steps. The top 10∼20 predicted medium combinations of the best objective values (e.g., *r*, *K*) were subjected to experimental validation. The results were included in the training data for the following round of ML model construction and prediction (Fig. 2B). Active learning was conducted for five rounds for each strain: R1 and R2 considering *r_Lp* or *K_Lp* for medium optimization of *Lp*, and S1∼S3 considering multiple parameters for the medium specialization of *Lp* or *Ec* (Fig. 2A). That is, S1-1 and S1-2 considered the pairs of *r* or *K* (i.e., *r_Lp* vs. *r_Ec*, *K_Lp* vs. *K_Ec*) to maximize the difference of *r* or *K* between *Lp* and *Ec*, and S2-1, S2-2, and S3 considered all parameters to maximize the difference of both *r* and *K* between *Lp* and *Ec* (Fig. 2C).

### Active learning successfully finetuned the media for selective bacterial growth

Active learning considering the single parameter of *r_Lp* and *K_Lp* (R1 and R2), which was started from the initial training data (R0), successfully increased *r_Lp* and *K_Lp* within two rounds; however, the media optimized for *Lp* also improved the growth of *Ec* (Fig. 3A-B, R1 and R2). Active learning considering multiple growth parameters was designed to maximize the difference of *r* or *K* between *Lp* and *Ec*, e.g., promoting the growth of *Lp* but repressing the growth of *Ec*. Three formulas were employed for the ML prediction and medium selection (see Methods). Three rounds of active learning considering multiple parameters) increased the medium specialization: significant *Lp* growth and no *Ec* growth (Fig. 3A-B, S1-1, S1-2, and S2-1). Intriguingly, although the optimization targeted a single parameter (*r_Lp* or *K_Lp*), the other was also improved to a certain extent (Fig. 3E-F, S1-1, S1-2, and S2-1). Moreover, the medium combinations suitable for *Ec* growth were successfully developed by active learning (Fig. 3C-D) despite the medium components originating from MRS, which is developed explicitly for *Lp*. Three rounds of active learning improved the growth of *Ec* but of poor specificity because *Lp* grew as well (Fig. 3C-D, S1-1, S1-2, and S2-1) and the parameters other than the targeted one were unsatisfied (Fig. 3G-H, S1-1, S1-2, and S2-1). Two additional rounds considerably increased the medium specialization for *Ec*, both the targeted (Fig. 3C-D, S2-2 and S3) and untargeted parameters (Fig. 3G-H, S2-2 and S3).

**Figure 3.**
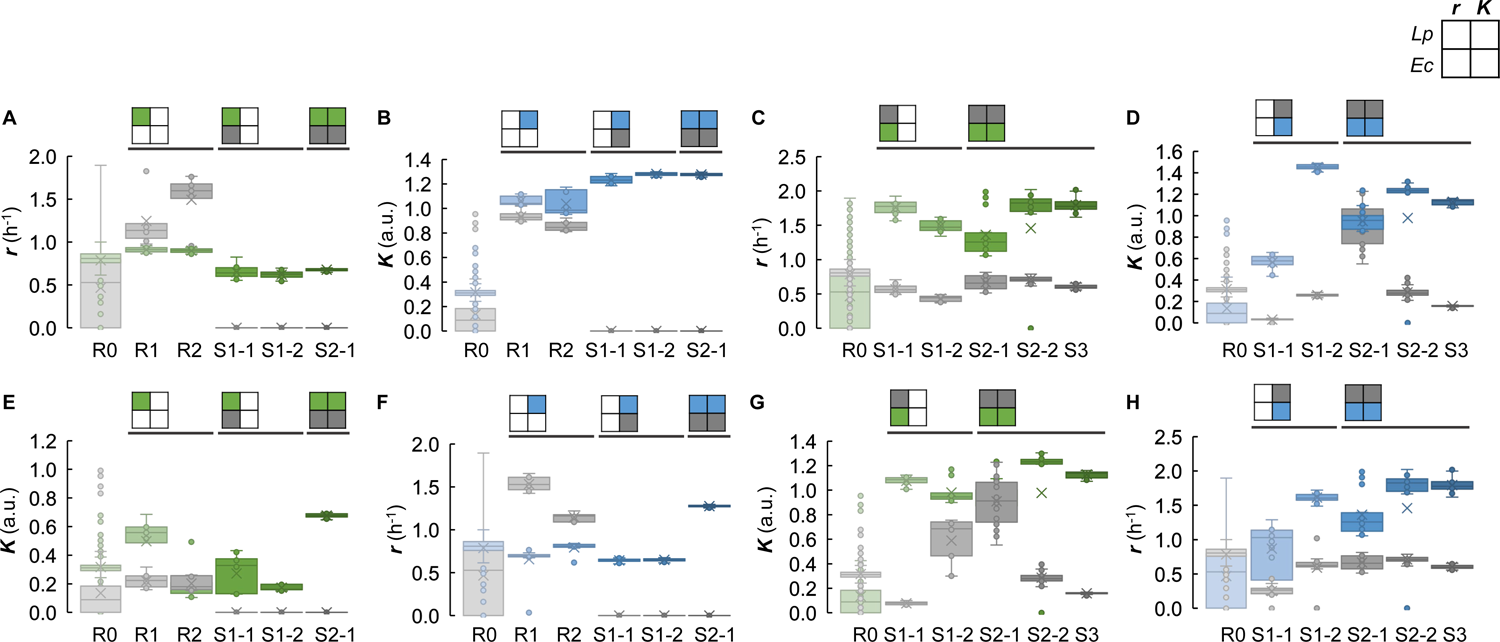
Active learning improved medium specialization. Medium optimization for the specific growth of *Lp* or *Ec* is shown as the left (**A**, **B**, **E**, **F**) and right (**C**, **D**, **G**, **H**) four panels, respectively. Boxplots represent the growth parameters experimentally obtained under 10∼20 medium combinations predicted per round of active learning. The colored boxes represent the parameters supposed to be improved in active learning, and the gray ones are to be repressed. The small quarter squares indicate the ML models used in active learning. The growth parameters to be improved in ML model construction are indicated in green and blue (*r* and *K*, respectively), and those to be suppressed are indicated in grey. R0 shows the initial training data. R1 and R2 show the experimental results of two rounds of active learning with the ML model considering either *r_Lp* or *K_Lp* solely. S1-1 and S1-2 show the experimental results of two rounds of active learning with the ML model considering the paired parameters of *r* or *K*. S2-1, S2-2, and S3 show the experimental results of three rounds of active learning with the ML models considering all four parameters.

Six medium combinations of high specificity for *Lp* (M1-3_Lp) or *Ec* (M1-3_Ec), newly developed via active learning, were selected for further verification. As the active learning prediction was performed in the mono-culture condition, whether these media remained specific in the presence of both *Lp* and *Ec* remained uncertain. The co-culture of *Lp* and *Ec* was performed in the six media, the medium compositions of which differed from that of MRS (Table 1). Nearly all of them exhibited significant specificity for the growth of the target strain, *Lp* or *Ec*, regardless of mono- or co-culture (Fig. 4). Although *Lp* producing acetic acid might inhibit *Ec* ^29, 30, 31^, the media developed for *Ec* growth (M1-3_Ec) remained specificity in the presence of *Lp*. The results suggested that the ML-assisted medium optimization and specialization were practical, and the resultant media were robust regardless of mono- or co-culture.

**Figure 4.**
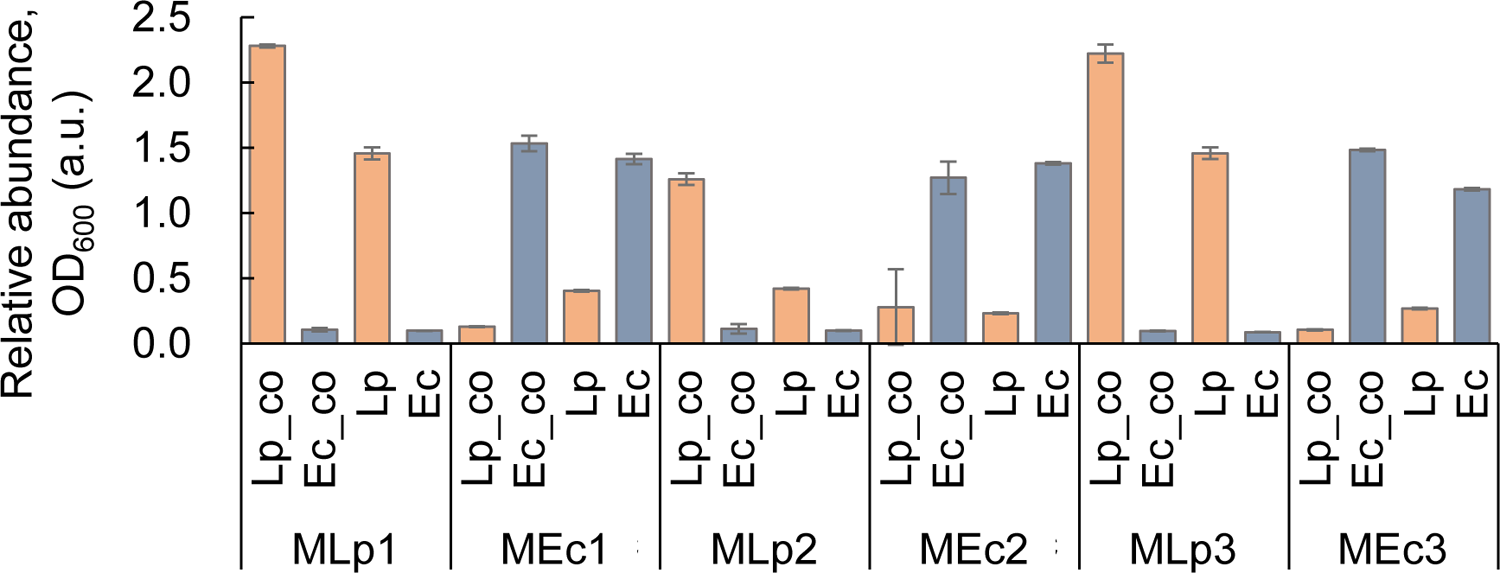
Co-culture of both strains in finetuned media for selective culture. Six media optimized for specific growth of either *Lp* or *Ec* are indicated as MLp1∼3 and MEc1∼3, respectively. The OD_600_ values of *Lp* and *Ec* after 24-h co-culture are shown. Orange and blue represent *Lp* and *Ec*, respectively. Standard errors of biological replicates (N=3) are indicated.

**Table 1.** ML predicted medium combinations of significant growth specificity. Six media were selected for the co-culture test. MLp1, 2, 3 and MEc1, 2, 3 indicate the ML predicted media specifically for *Lp* and *Ec* growth, respectively. The concentrations of the medium components are shown in the unit of g/L in comparison to the original medium MRS.

| (g/L) | MLp1 | MEc1 | MLp2 | MEc2 | MLp3 | MEc3 | MRS |
| --- | --- | --- | --- | --- | --- | --- | --- |
| Tryptone | 10 | 10 | 10 | 10 | 10 | 10 | 10 |
| Yeast extract | 5 | 1.35 | 1.35 | 0.05 | 5 | 1.35 | 5 |
| Glucose | 16.2 | 0.6 | 0.6 | 0.2 | 16.2 | 0.2 | 20 |
| K<sub>2</sub>HPO<sub>4</sub> | 0.18 | 4.86 | 6 | 6 | 0.18 | 6 | 6 |
| C<sub>6</sub>H<sub>14</sub>N<sub>2</sub>O<sub>7</sub> | 1.62 | 0.54 | 0.54 | 0.18 | 1.62 | 0.18 | 2 |
| CH<sub>3</sub>COONa | 4.05 | 0.15 | 0.15 | 0.15 | 4.05 | 0.15 | 15 |
| MgSO<sub>4</sub> | 0.6 | 0.6 | 0.486 | 0.006 | 0.6 | 0.018 | 0.6 |
| MnSO<sub>4</sub> | 0.16 | 0.1296 | 0.1296 | 0.0016 | 0.16 | 0.0144 | 0.16 |
| FeSO<sub>4</sub> | 0.0005 | 0.0045 | 0.0045 | 0.0135 | 0.0005 | 0.0135 | 0.05 |
| CH<sub>3</sub>COOH | 0.013 | 0.013 | 3.159 | 0.013 | 0.013 | 0.039 | 1.3 |
| Tween80 | 1 | 0.27 | 0.27 | 0.81 | 1 | 0.81 | 1 |

### Changes in growth parameters revealed the effectiveness of active learning

How the growth was finetuned during the active learning was further analyzed. The distributions of the growth parameters significantly differentiated between the strains, considerably changing during active learning (Fig. 5A). Bimodal distributions were commonly observed in *Ec* (Fig. 5A, bottom), indicating the medium combinations predicted in the active learning were highly selective for *Ec* growth. In contrast, monomodal distributions were more often found in *Lp*, although the transition from monomodal to bimodal was observed in *r_Lp* (Fig. 5A, upper). The peaks of distributions altered significantly along with the active learning proceeded (Fig. 5, color variation), revealing the effectiveness of active learning for medium optimization and specialization. In addition, correlation analysis of the four growth parameters showed significant cross-correlations, except for the pair of *r_Ec* and *r_Lp* (Fig. 5B, red). The positive correlations between *r* and *K* in both strains (Fig. 5B, blue) indicated a common feature of improved growth rate associated with increased population density. The negative correlation across the strains (Fig. 5B, black) reasonably presented the trade-offs in the growth of *Lp* and *Ec*, as the active learning aimed to improve the medium specificity for a single strain. Taken together, the features of the datasets acquired during active learning well reflected the process of medium optimization and specialization.

**Figure 5.**
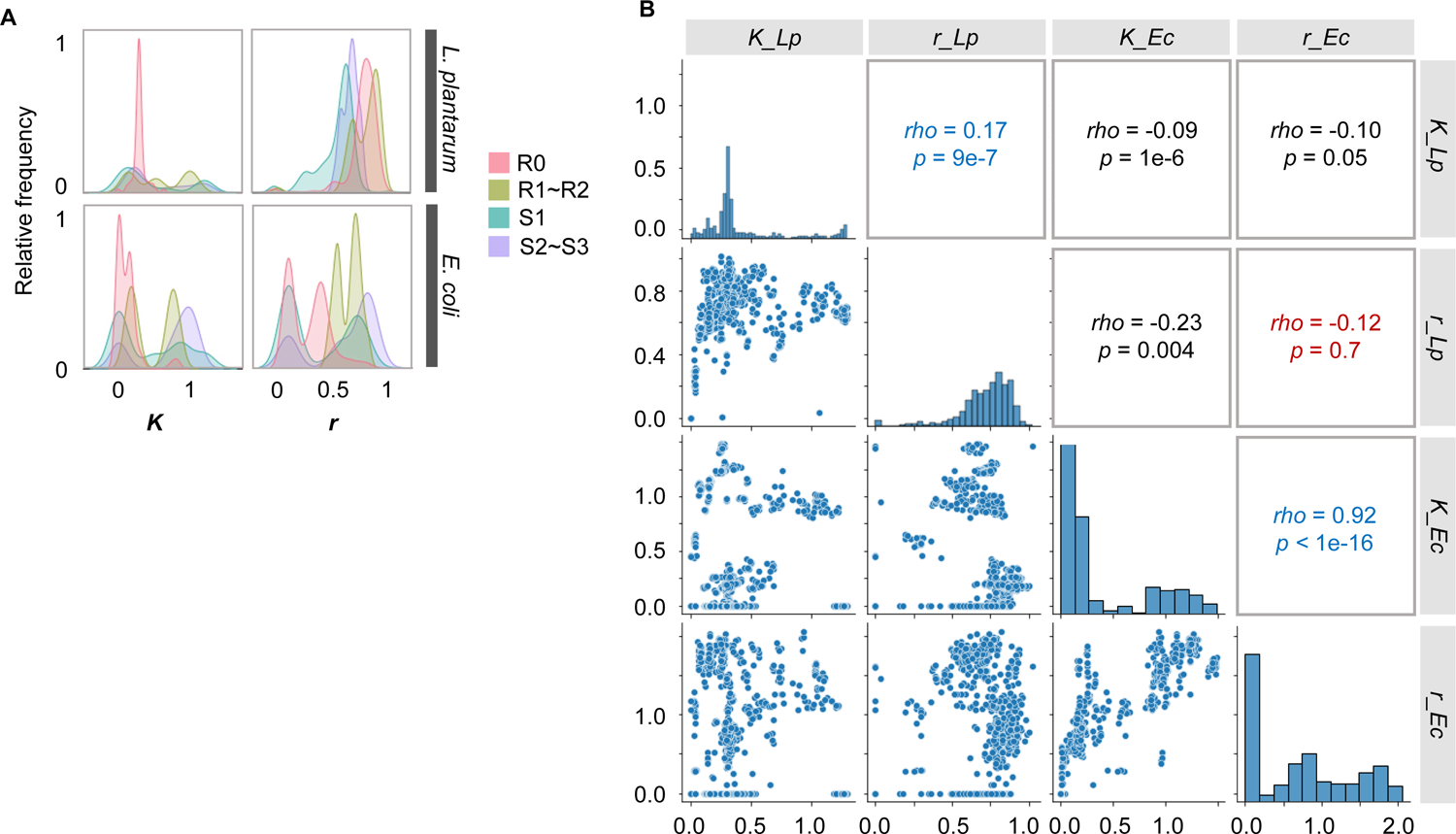
Data analysis of the growth parameters acquired in active learning. **A.** Distributions of the growth parameters. The relative frequency of the parameters acquired in the indicated rounds of active learning is shown. Color variation indicates the rounds of active learning. **B.** Correlation analysis of the four growth parameters. Scatter plots, histograms, correlation coefficients, and statistical significance are presented.

### Decision-making medium components for the changes in bacterial growth

GBDT analysis showed that all four growth parameters were primarily determined by a single medium component (Fig. 6). Differentiated decision-making components were observed in *Lp*, i.e., yeast extract and acetic acid for *K* and *r*, respectively (Fig. 6, upper). As yeast extract was reported to provide initial nutrients for cell division and substance synthesis ^32, 33^, the abundance of the resource might determine the final population size. It was reasonable that acetic acid, which often inhibited microbe growth ^34, 35, 36^, decided *r_Lp*, as Lp preferred an acidic environment. On the other hand, both *r_Ec* and *K_Ec* were commonly determined by K_2_HPO_4_ (Fig. 6, bottom), which might provide the buffering effect in response to the changes in pH caused by *Lp*.

**Figure 6.**
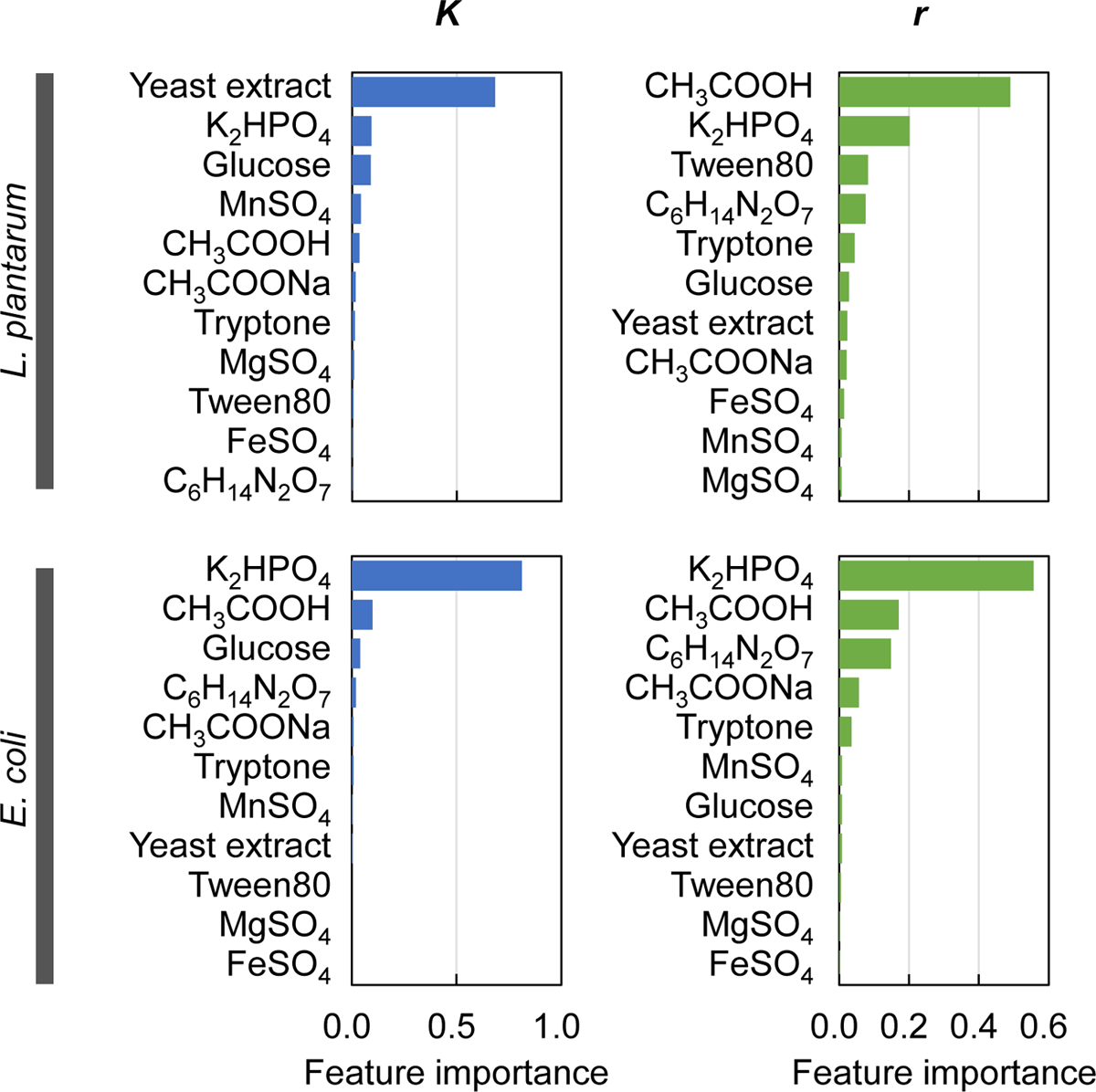
Contribution of medium components to bacterial growth. Feature importance of the medium components analyzed with GBDT is shown. Blue and green represent the growth yield (*K*) and the growth rate (*r*), respectively. The bacterial strains are indicated.

Hierarchical clustering analysis of the normalized feature importance intriguingly divided the medium components into four main categories (Fig. 7A). The medium components assigned in the same categories showed neither common chemical properties nor similar biological functions. It strongly suggested that the novel classification of medium components depended on their impact on bacterial growth. Four different trends of medium components contributing to the growth parameters were identified, that is, highly relevant to *K_Lp* (pink), *r_Lp* (yellow), *r* (purple), and *Ec* (grey), respectively (Fig. 7B). Such specificity of medium components might be applied to the medium development for differentiated bacterial growth.

**Figure 7.**
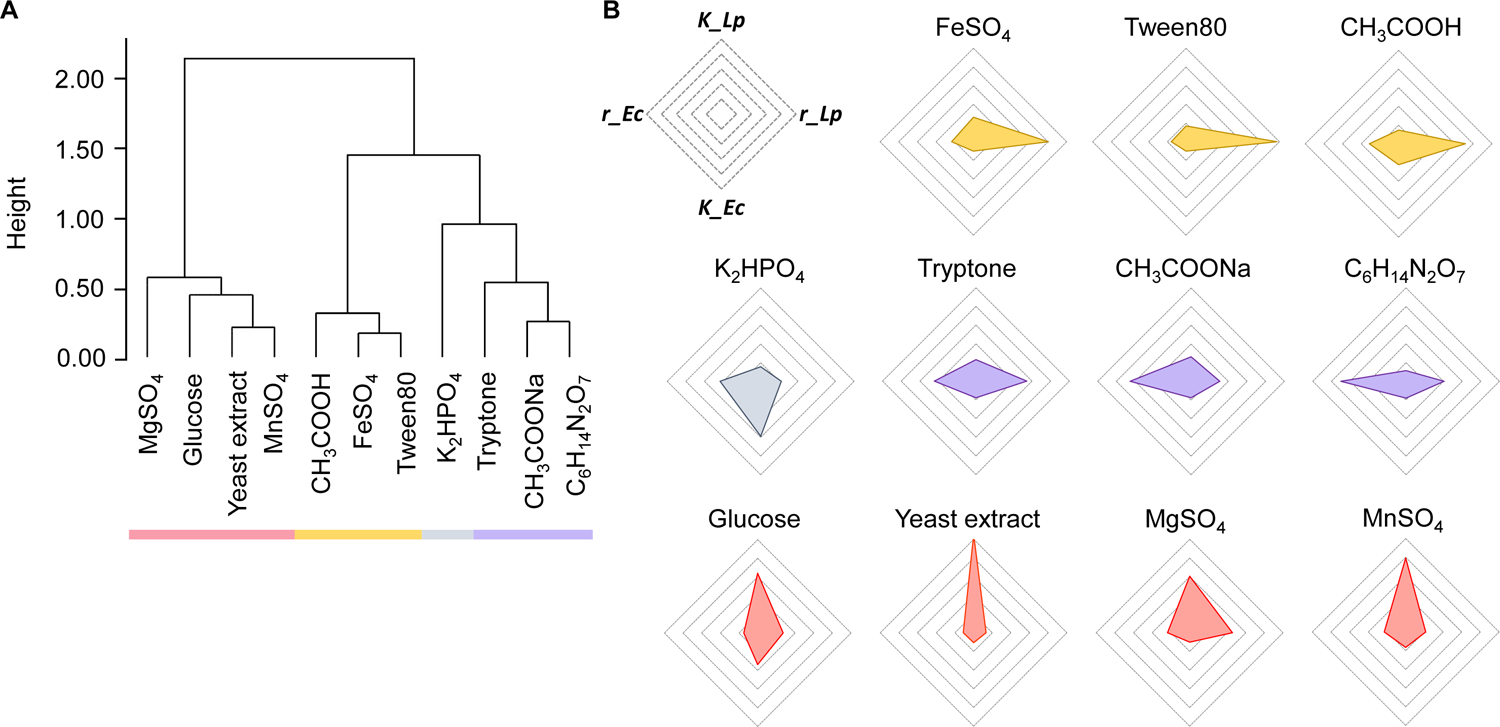
Categorization of medium components according to their contributions to bacterial growth. **A.** Hierarchical clustering of medium components. **B.** Radar plots of the feature importance of medium components to the four growth parameters. Color variation indicates the four categories.

### Medium components adjusted via active learning for bacterial growth specificity

The medium components contributing to the bacterial growth specificity could be identified according to the medium specialization proceeded *via* active learning. The scores (*S*), calculated in active learning considering multiple growth parameters, were subjected to GBDT analysis. As they represented the goodness of the growth specificity, the medium components of high feature importance indicated a significant contribution to growth specificity. The results showed that yeast extract and glucose primarily determined the specificity of *K* for *Lp* and *Ec*, respectively (Fig. 8A, blue), and K_2_HPO_4_ was the common component determining the specificity of *r* for *Lp* and *Ec* (Fig. 8A, green). yeast extract and K_2_HPO_4_ (Fig. 8A, black). The findings revealed that adjusting a single component differentiated the growth of *Lp* and *Ec* to a great extent.

**Figure 8.**
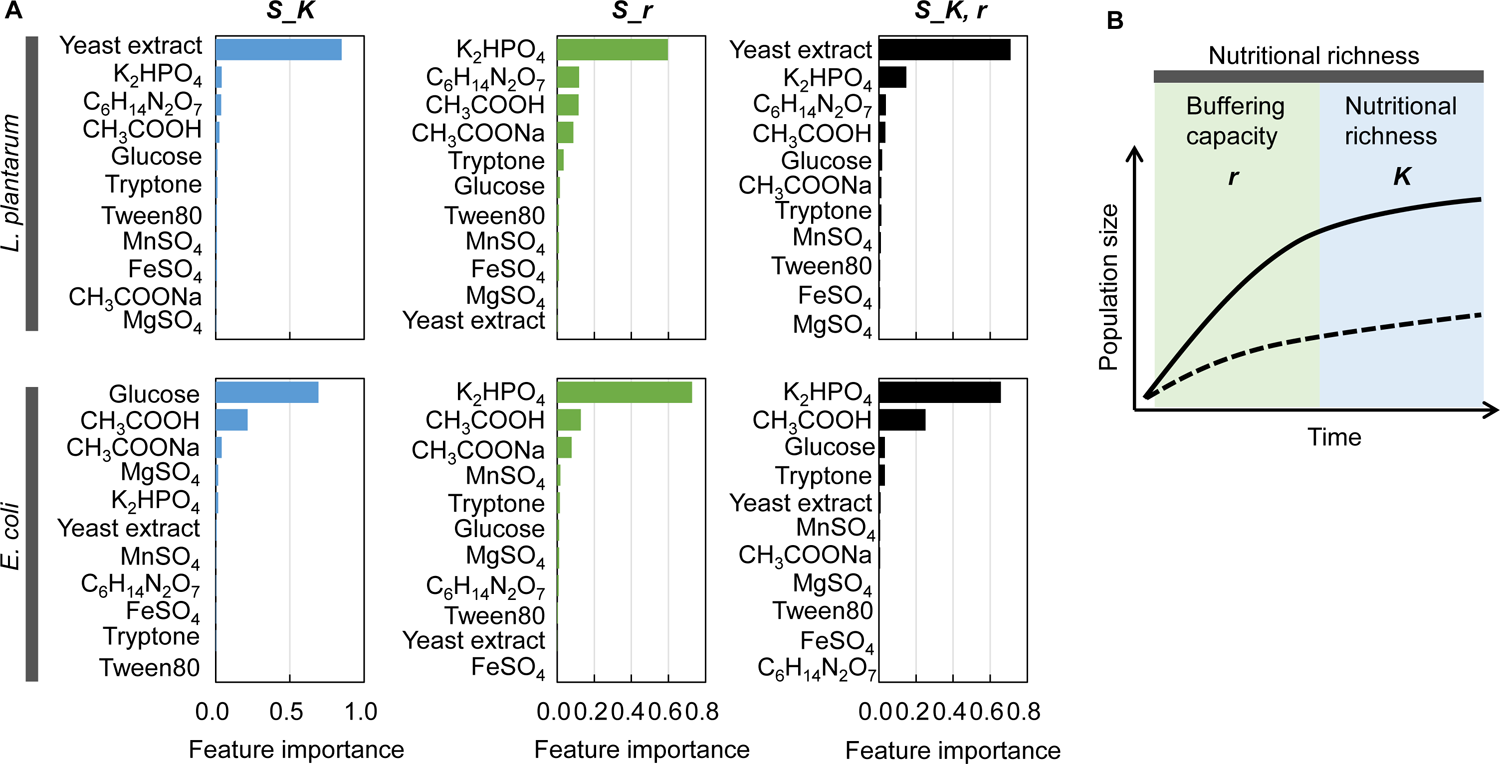
Contribution of medium components to the growth differentiation. **A.** Feature importance of medium components to the scores considering multiple growth parameters. *S_K*, *S_r*, *S_K,* and *r* indicate the scores considering the pair of *K_Lp* and *K_Ec*, the pair of *r_Lp* and *r_Ec*, and all four parameters, respectively. **B.** Hypothesis of differentiated factors determining the bacterial growth specificity in exponential and stationary phases.

In summary, the bacterial growth specificity was roughly determined by a single component, regardless of considering single or both parameters of *r* and *K*. As *r* and *K* were the most representative features that quantitatively described bacterial growth dynamics ^37, 38^, the determinative manner of medium components contributing to *r* and *K* revealed the working principle for specific growth control. Buffering capacity and nutritional richness seemed to influence the growth specificity during the exponential and stationary phases, respectively (Fig. 8B). K_2_HPO_4_ was supposed to control the pH condition and execute the buffering effect. Yeast extract and glucose were considered to provide the nutrients, such as carbon resources, for metabolism supporting bacterial growth. The differentiation in the essential components for bacterial growth specificity was well supported by the findings that the buffering agents influenced cell division and biosynthesis ^39, 40^ and the nutritional resources affected the organic acid metabolism ^41, 42^. In summary, ML-assisted medium optimization and specialization provided a practical tool for medium development and discovered novel insights into bacterial growth for precise culture control.

## Discussion

The present study first demonstrated ML-assisted medium specialization for differentiated bacterial growth. ML was remarkably significant in medium optimization for microbial and mammalian cells ^21, 26^, which could be widely applied to synthetic construction and production ^43, 44^. The present study provided alternative applications in medium development for selective culture. The results indicated that combining ML with active learning was highly practical for precise finetuning of the medium compositions for the selective culture of particular bacteria. Further applications of optimizing selective culture media for complex microbiomes were perceived, such as the systematic development of selective media for individual bacteria living in the environmental microbial community.

The present study made a first trial to combine the growth parameters, determined according to the experimental records, as the quantitative reference values for model construction and prediction. Medium optimization for multiple strains might raise the cost of data acquisition for training and testing. Theoretically, more data led to a more accurate ML model, and more targets (variables) required more experimental data ^45^. To save labor and cost, three combinations of four growth parameters (*r_Lp*, *K_Lp*, *r_Ec*, and *K_Ec*), representing the growth features of two different bacterial strains, were employed in the active learning here. The success in medium optimization demonstrated that combining multiple parameters was highly recommended for finetuning the selective culture media. Note that many other combinations of the growth parameters could be considered, which might show higher efficiency or better selectivity.

On the other hand, improving multiple growth parameters simultaneously in model construction was theoretically ideal but might be biologically impractical, as the living cells and their communities were highly self-regulated and coordinated ^46, 47, 48, 49^. In the present study, the constructed ML models tried to improve the growth rate (*r*) and maximize the population size (*K*) simultaneously, which was assumed to be impossible because of the potential trade-offs between *r* and *K* ^50, 51^. Intriguingly, active learning allowed us to find the medium combinations that improve both *r* and *K*, demonstrating the availability of the parallel optimization of multiple growth parameters. The differentiated growth of two bacterial strains was also successfully achieved when considering the growth parameters of both strains. Intriguingly, the selective culture media developed in the mono-culture maintained the specificity for bacterial growth in the co-culture. As the two strains (*Lp* and *Ec*) in the present study were ecologically and genetically far from each other, whether the present approach for medium specialization was practical for closely related or habited bacteria remained questioned. If single species played the dominant role in the microbial communities ^52, 53, 54, 55^, the interactions among multiple species might be ignored in the active learning by weighting the particular growth parameters in ML models. Nevertheless, further technical and experimental developments are required.

In addition, the big dataset acquired in active learning allowed us to investigate the contribution of medium components to bacterial growth. A novel understanding of the chemical role of bacterial growth was achieved. As an example of new findings, acetic acid was commonly used to adjust the pH of the media for culturing *Lp* (e.g., MRS) to suppress the growth of other microbiomes growing at neutral conditions ^34, 35, 36^. The GBDT analytical results showed that the inhibitory effect of acetic acid on *Ec* was limited, and yeast extract played a more significant role in selective culture. The finding indicated that the commonly used or commercially available media could be further finetuned for better performance or milder conditions. For instance, antibiotics or dye were often added to the media for selective microbial culture, which might cause increased resistance due to frequent usage ^56, 57, 58, 59^. Optimizing the medium components other than antibiotics should be challenged to acquire milder conditions for suppressed bacterial growth associated with reduced potentiality of acquiring antibiotic resistance.

ML-assisted medium optimization often resulted in novel insights that were out of the knowledge. Besides the present findings of the medium contributions to differentiated bacterial growth, our previous studies observed the secondary contribution of glucose to bacterial growth ^60^, the differentiated contribution of carbon, sulfate, and nitrogen for survival ^27^ and the diversified metabolic strategies in transcriptome reorganization for increased productivity ^26^. Additionally, the cluster analysis intriguingly divided the medium components into four clusters, which were out of any well-known chemical or biological categories. The mono-culture data showed higher accuracy in predicting interspecies relationships than the metabolic or phylogenetic data ^28^. Taken together, active learning for medium optimization and specialization allowed better cell culture and provided the dataset connecting medium compositions to microbial growth for better understanding and application of microorganisms.

## Materials and Methods

### Bacterial strains

*Escherichia coli* BW25113 and *Lactobacillus plantarum* (ATCC8014) were used, which were obtained from the National BioResource Project, National Institute of Genetics (Shizuoka, Japan), and the National Biological Resource Center (Tsukuba, Japan), respectively. As previously described in detail ^60, 61^, the stock solutions of the bacterial cells grown in the exponential phase were prepared for growth assay in advance, and hundreds of the stock solutions (60 µL) were stored at −80°C for future use.

### Medium combinations

Medium combinations were initially decided according to the commercially available medium MRS (Wako). The components (chemical compounds, reagents, etc.) comprised in MRS were purchased from Wako, except Tryptone (Sigma), yeast extract (MP Biochemicals), and Tween 80 (MP Biochemicals). The lowest concentrations of these components were set at 1% of their concentrations in MRS. Their highest concentrations were determined based on the literature and manufacturers’ instructions. The stock solutions of the medium components were prepared as described previously ^27, 60^. They were aliquoted into 100∼1,000 µL portions for single use and stored at −30°C. The medium combinations were prepared by mixing the stock solutions, of which the concentrations varied logarithmically in five different gradients, as previously reported ^26, 27, 60^. Initially, 96 medium combinations were prepared for the growth assay of both bacterial strains as the training data. A total of 192 combinations were prepared to test both strains in the present study.

### Growth assay and calculation

The prepared culture mixtures were dispensed into a 96-well microplate (Costar) with 3-4 biological replicates per combination, each consisting of 200 μL per well, and the combinations were placed in different positions. The 96-well plate was incubated at 37°C with shaking at 567 rpm in a microplate reader (Epoch2, BioTek). Cell growth was monitored at an optical density of 600 nm (OD_600_), and readings were taken at 30 min intervals over 48 h. The temporal changes of OD_600_ readings were exported from the microplate reader and subjected to the Python programs to calculate the two representative growth parameters, the growth rate (*r*) and the maximal OD_600_ (*K*), as described elsewhere in detail^60, 61^. In total, 1,660 growth curves were experimentally obtained, and the mean values of *r* and *K* (biological replicates, N=4∼6) were used for machine learning and computational analyses.

### Machine learning and computational analyses

Python was used for machine learning (ML), as described previously ^21, 26, 27^. The ML models used “GradientBoostingRegressor” from the “ensemble” module in the “scikit-learn” library. The explanatory and target variables were the medium components and growth parameters. The model employed ‘random_state’ and ‘n_estimators’ set to 0 and 300, respectively. The ‘learning_rate’ and ‘max_depth’ were searched between 0.001 and 0.5, using increments of 0.005 among 2, 3, 4, and 5. Root mean square error (RMSE) was calculated to assess prediction accuracy, using the ‘mean_squared_error’ from the ‘metrics’ module in “scikit-learn”. The ‘feature_importances_’ was calculated using an outer five-fold cross-validation. Both outer and inner cross-validations were performed using the ‘cross_val_score’ function from the ‘model_selection’ module in “scikit-learn”. ‘GridSearchCV’ was used for hyperparameter search with ‘learning_rate’ and ‘max_depth’ ranging between 0.01 and 0.5, incrementing by 0.01 among 2, 3, 4, and 5. The ‘n_estimators’ were set to 300, while other hyperparameters were left as default. The average of the ‘feature_importances_’ values derived from the five-fold cross-validation was used. Additionally, the ‘feature_importances_’ were subjected to the hierarchical clustering analysis, using ‘normalize’ in the “sklearn.preprocessing” package with the “ ward “ method.

### Model construction and active learning

As previously reported ^21^, ML model construction and prediction were conducted using the supercomputer Cygnus system (NEC LX 124Rh-4G) in active learning. The GBDT models (R0∼R2) were constructed initially in active learning, i.e., learning, prediction, and validation. The top 10∼20 predicted medium combinations that showed the best *r* or *K* of individual strain were subjected to experimental verification. The resultant experimental outputs were included in the training dataset, which was used for the following round of ML model construction. Subsequently, the ML models combining the multiple growth parameters (*r* and *K*) were constructed using the following formula (Eqs. 1∼3) in the following rounds of active learning.

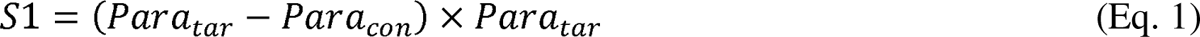

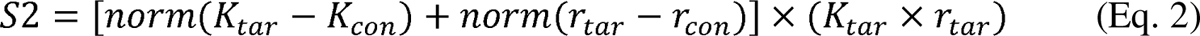

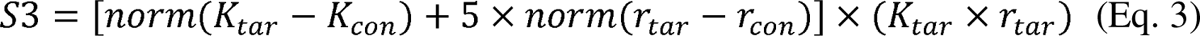

Here, *Para* represents any of the growth parameters of any strain. *Para_tar* and *Para_con* indicated the parameters of the target and control strain, respectively. *Norm* indicated the data normalization. *K_tar* and *K_con*, *r_tar*, and *r_con* represented *K* and *r* as the target or control, respectively. The resultant scores (*S1*∼*S3*) were used as the target variables. The higher the scores, the more significant the difference in growth parameters, indicating higher specificity for the target bacterial growth. The top 10∼20 medium combinations showing the highest scores (*S1*∼*S3*) were experimentally tested and to the training dataset for subsequent learning and prediction. Repeated rounds of active learning associated with the changes in ML models were performed.

### Co-culture verification

The cell stocks of both bacterial strains were diluted 1,000-fold in 1 mL of the identical medium separately. The diluted mixture of *E. coli* was dispensed into a 24-well plate, and that of *L. plantarum* was placed into a Transwell insert (ThinCert® Cell Culture Inserts, pore size 0.4 µm, Greiner Bio-One) and then positioned on the 24-well plate (Greiner Bio-One) containing the *E. coli* mixture. It allowed the two bacterial strains to grow in the same medium condition without mixing each other. The 24-well plate was incubated at 37°C with shaking at 567 rpm (Epoch2, BioTek) for 24 h. The culture mixtures were individually injected into separate wells in an alternative 24-well microplate (1 mL per well) and subjected to measure the OD_600_ readings using the same microplate reader. Six finetuned selective media were tested with three biological replicates.

## Acknowledgments

This work was partially supported by the JSPS KAKENHI Grant-in-Aid for Challenging Exploratory Research (21K19815) and by the JSPS KAKENHI Grant-in-Aid for Scientific Research (B) (19H03215).

## References

1. Hugon P, Dufour JC, Colson P, Fournier PE, Sallah K, Raoult D. A comprehensive repertoire of prokaryotic species identified in human beings. Lancet Infect Dis 15, 1211–1219 (2015).

2. Lagier JC, et al. Microbial culturomics: paradigm shift in the human gut microbiome study. Clin Microbiol Infect 18, 1185–1193 (2012).

3. Lagier JC, et al. Culturing the human microbiota and culturomics. Nat Rev Microbiol 16, 540–550 (2018).

4. Bonnet M, Lagier JC, Raoult D, Khelaifia S. Bacterial culture through selective and non-selective conditions: the evolution of culture media in clinical microbiology. New Microbes New Infect 34, 100622 (2020).

5. Rappé MS, Connon SA, Vergin KL, Giovannoni SJ. Cultivation of the ubiquitous SAR11 marine bacterioplankton clade. Nature 418, 630–633 (2002).

6. Prasad M, Shetty SK, Nair BG, Pal S, Madhavan A. A novel and improved selective media for the isolation and enumeration of Klebsiella species. Appl Microbiol Biotechnol 106, 8273–8284 (2022).

7. Inoue Y. Three semi-selective media for Pseudomonas syringae pv. maculicola and P. cannabina pv. alisalensis. Appl Microbiol Biotechnol 106, 5741–5755 (2022).

8. Cook BS, Beddow JG, Manso-Silván L, Maglennon GA, Rycroft AN. Selective medium for culture of Mycoplasma hyopneumoniae. Vet Microbiol 195, 158–164 (2016).

9. Chon JW, Hyeon JY, Park JH, Song KY, Kim JH, Seo KH. Improvement of mannitol-yolk-polymyxin B agar by supplementing with trimethoprim for quantitative detection of Bacillus cereus in foods. J Food Prot 75, 1342–1345 (2012).

10. Kim DH, Kim H, Chon JW, Moon JS, Song KY, Seo KH. Development of blood-yolk-polymyxin B-trimethoprim agar for the enumeration of Bacillus cereus in various foods. Int J Food Microbiol 165, 144–147 (2013).

11. Azubuike CC, Edwards MG, Gatehouse AMR, Howard TP. Applying Statistical Design of Experiments To Understanding the Effect of Growth Medium Components on Cupriavidus necator H16 Growth. Appl Environ Microbiol 86, (2020).

12. Singh V, Haque S, Niwas R, Srivastava A, Pasupuleti M, Tripathi CK. Strategies for Fermentation Medium Optimization: An In-Depth Review. Front Microbiol 7, 2087 (2016).

13. Hemalatha M, Subathra Devi C. A statistical optimization by response surface methodology for the enhanced production of riboflavin from Lactobacillus plantarum-HDS27: A strain isolated from bovine milk. Front Microbiol 13, 982260 (2022).

14. Nettoor Veettil V, A VC. Optimization of bacteriocin production by Lactobacillus plantarum using Response Surface Methodology. Cell Mol Biol (Noisy-le-grand) 68, 105–110 (2022).

15. Singh V, Khan M, Khan S, Tripathi CK. Optimization of actinomycin V production by Streptomyces triostinicus using artificial neural network and genetic algorithm. Appl Microbiol Biotechnol 82, 379–385 (2009).

16. Camacho DM, Collins KM, Powers RK, Costello JC, Collins JJ. Next-Generation Machine Learning for Biological Networks. Cell 173, 1581–1592 (2018).

17. Vamathevan J, et al. Applications of machine learning in drug discovery and development. Nat Rev Drug Discov 18, 463–477 (2019).

18. AlQuraishi M. Machine learning in protein structure prediction. Curr Opin Chem Biol 65, 1–8 (2021).

19. Gao Y, et al. Machine learning based early warning system enables accurate mortality risk prediction for COVID-19. Nat Commun 11, 5033 (2020).

20. Cosenza Z, Block DE, Baar K. Optimization of muscle cell culture media using nonlinear design of experiments. Biotechnol J 16, e2100228 (2021).

21. Hashizume T, Ozawa Y, Ying BW. Employing active learning in the optimization of culture medium for mammalian cells. NPJ Syst Biol Appl 9, 20 (2023).

22. Jang J, Hur HG, Sadowsky MJ, Byappanahalli MN, Yan T, Ishii S. Environmental Escherichia coli: ecology and public health implications-a review. J Appl Microbiol 123, 570–581 (2017).

23. Huleani S, Roberts MR, Beales L, Papaioannou EH. Escherichia coli as an antibody expression host for the production of diagnostic proteins: significance and expression. Crit Rev Biotechnol 42, 756–773 (2022).

24. Colautti A, Orecchia E, Comi G, Iacumin L. Lactobacilli, a Weapon to Counteract Pathogens through the Inhibition of Their Virulence Factors. J Bacteriol 204, e0027222 (2022).

25. van de Wijgert J, Verwijs MC. Lactobacilli-containing vaginal probiotics to cure or prevent bacterial or fungal vaginal dysbiosis: a systematic review and recommendations for future trial designs. Bjog 127, 287–299 (2020).

26. Aida H, et al. Machine learning-assisted medium optimization revealed the discriminated strategies for improved production of the foreign and native metabolites. Comput Struct Biotechnol J 21, 2654–2663 (2023).

27. Aida H, Hashizume T, Ashino K, Ying BW. Machine learning-assisted discovery of growth decision elements by relating bacterial population dynamics to environmental diversity. Elife 11, (2022).

28. Nestor E, Toledano G, Friedman J. Interactions between Culturable Bacteria Are Predicted by Individual Species’ Growth. mSystems 8, e0083622 (2023).

29. Aditya A, Rahaman SO, Biswas D. Impact of Lactobacillus-originated metabolites on enterohemorrhagic E. coli in rumen fluid. FEMS Microbiol Ecol 98, (2022).

30. Yan R, et al. Anticolonization of Carbapenem-Resistant Klebsiella pneumoniae by Lactobacillus plantarum LP1812 Through Accumulated Acetic Acid in Mice Intestinal. Front Cell Infect Microbiol 11, 804253 (2021).

31. Mukherjee A, Ealy J, Huang Y, Benites NC, Polk M, Basan M. Coexisting ecotypes in long-term evolution emerged from interacting trade-offs. Nat Commun 14, 3805 (2023).

32. Tao Z, et al. Yeast Extract: Characteristics, Production, Applications and Future Perspectives. J Microbiol Biotechnol 33, 151–166 (2023).

33. Jiang M, Chen K, Liu Z, Wei P, Ying H, Chang H. Succinic acid production by Actinobacillus succinogenes using spent brewer’s yeast hydrolysate as a nitrogen source. Appl Biochem Biotechnol 160, 244–254 (2010).

34. Fraise AP, Wilkinson MA, Bradley CR, Oppenheim B, Moiemen N. The antibacterial activity and stability of acetic acid. J Hosp Infect 84, 329–331 (2013).

35. Ge J, Kang J, Ping W. Effect of Acetic Acid on Bacteriocin Production by Gram-Positive Bacteria. J Microbiol Biotechnol 29, 1341–1348 (2019).

36. Feng L, et al. Evaluation of the antibacterial, antibiofilm, and anti-virulence effects of acetic acid and the related mechanisms on colistin-resistant Pseudomonas aeruginosa. BMC Microbiol 22, 306 (2022).

37. Blomberg A. Measuring growth rate in high-throughput growth phenotyping. Curr Opin Biotechnol 22, 94–102 (2011).

38. Peleg M, Corradini MG. Microbial growth curves: what the models tell us and what they cannot. Crit Rev Food Sci Nutr 51, 917–945 (2011).

39. Jingjing E, et al. Effects of buffer salts on the freeze-drying survival rate of Lactobacillus plantarum LIP-1 based on transcriptome and proteome analyses. Food Chem 326, 126849 (2020).

40. Jingjing E, et al. Improving the freeze-drying survival rate of Lactobacillus plantarum LIP-1 by increasing biofilm formation based on adjusting the composition of buffer salts in medium. Food Chem 338, 128134 (2021).

41. De Bruyn IN, Holzapfel WH, Visser L, Louw AI. Glucose metabolism by Lactobacillus divergens. J Gen Microbiol 134, 2103–2109 (1988).

42. Mogodiniyai Kasmaei K, Schlosser D, Sträuber H, Kleinsteuber S. Does glucose affect the de-esterification of methyl ferulate by Lactobacillus buchneri? Microbiologyopen 9, e971 (2020).

43. Aida H, Ying BW. Efforts to Minimise the Bacterial Genome as a Free-Living Growing System. Biology (Basel) 12, (2023).

44. Radivojevic T, Costello Z, Workman K, Garcia Martin H. A machine learning Automated Recommendation Tool for synthetic biology. Nat Commun 11, 4879 (2020).

45. Settles B. Active learning literature survey.). Computer Sciences Technical Report 1648, University of Wisconsin-Madison (2009).

46. Vargas-Garcia CA, Ghusinga KR, Singh A. Cell size control and gene expression homeostasis in single-cells. Curr Opin Syst Biol 8, 109–116 (2018).

47. Orland C, Emilson EJS, Basiliko N, Mykytczuk NCS, Gunn JM, Tanentzap AJ. Microbiome functioning depends on individual and interactive effects of the environment and community structure. ISME J, (2018).

48. Nishimura I, Kurokawa M, Liu L, Ying BW. Coordinated Changes in Mutation and Growth Rates Induced by Genome Reduction. mBio 8, (2017).

49. Rivett DW, et al. Resource-dependent attenuation of species interactions during bacterial succession. ISME J 10, 2259–2268 (2016).

50. Engen S, Saether BE. r- and K-selection in fluctuating populations is determined by the evolutionary trade-off between two fitness measures: Growth rate and lifetime reproductive success. Evolution 71, 167–173 (2017).

51. Luckinbill LS. r and K Selection in Experimental Populations of Escherichia coli. Science 202, 1201–1203 (1978).

52. Baichman-Kass A, Song T, Friedman J. Competitive interactions between culturable bacteria are highly non-additive. Elife 12, (2023).

53. Gould AL, et al. Microbiome interactions shape host fitness. Proc Natl Acad Sci U S A 115, E11951–e11960 (2018).

54. Piccardi P, Vessman B, Mitri S. Toxicity drives facilitation between 4 bacterial species. Proc Natl Acad Sci U S A 116, 15979–15984 (2019).

55. Palmer JD, Foster KR. Bacterial species rarely work together. Science 376, 581–582 (2022).

56. Ma Y, Ramoneda J, Johnson DR. Timing of antibiotic administration determines the spread of plasmid-encoded antibiotic resistance during microbial range expansion. Nat Commun 14, 3530 (2023).

57. Nordholt N, Kanaris O, Schmidt SBI, Schreiber F. Persistence against benzalkonium chloride promotes rapid evolution of tolerance during periodic disinfection. Nat Commun 12, 6792 (2021).

58. Pereira C, Warsi OM, Andersson DI. Pervasive Selection for Clinically Relevant Resistance and Media Adaptive Mutations at Very Low Antibiotic Concentrations. Mol Biol Evol 40, (2023).

59. Zheng EJ, et al. Modulating the evolutionary trajectory of tolerance using antibiotics with different metabolic dependencies. Nat Commun 13, 2525 (2022).

60. Ashino K, Sugano K, Amagasa T, Ying BW. Predicting the decision making chemicals used for bacterial growth. Sci Rep 9, 7251 (2019).

61. Kurokawa M, Ying BW. Precise, High-throughput Analysis of Bacterial Growth. J Vis Exp, (2017).

